# Environmental control of morphological color changes in the Bigfin Reef Squid, Sepioteuthis lessoniana

**DOI:** 10.64898/2026.04.23.720393

**Authors:** A. Chang, D.E. Kim, A. Hoffman, Z. Lajbner, J. Miller, S.G. Alvarado

## Abstract

Phenotypic plasticity enables organisms to adjust their traits in response to environmental conditions, yet studies of animal coloration—particularly in cephalopods—have largely focused on rapid neural control while overlooking slower, substrate-level changes. Here, we investigate whether morphological color change contributes to phenotypic plasticity in the bigfin reef squid (*Sepioteuthis lessoniana*). Hatchlings were reared in contrasting light (white) and dark (black) environments for 14 weeks, and chromatophore organization was quantified across developmental stages (4, 9, and 14 weeks) and body regions (dorsal and ventral) using high-resolution imaging and automated analysis. We find that environmental context and age interact to shape both the density and size of black, yellow, and red chromatophores, revealing a consistent trade-off between chromatophore number and expansion. Black-reared individuals maintain higher chromatophore densities, whereas white-reared individuals exhibit fewer but larger cells, particularly at later developmental stages. These effects are region-specific and align with the emergence of countershading, suggesting ecological tuning of pigmentation architecture. Our results demonstrate that the cellular substrate underlying color change is itself plastic and dynamically remodeled across development. We propose a hierarchical model in which slow morphological plasticity defines the functional limits of rapid neural color change. This work expands current frameworks of cephalopod coloration by integrating developmental, environmental, and cellular mechanisms, and highlights the importance of multi-timescale plasticity in adaptive visual signaling.

## INTRODUCTION

Phenotypic plasticity, the ability of an organism to produce a range of relatively adaptive phenotypes in response to dynamic environmental changes, is a fundamental mechanism of survival and adaptation (DeWitt et al. 1998). Plasticity can manifest at multiple levels of biological organization. At the morphological level, organisms alter their physical form in response to environmental cues, as in *Daphnia*, which develops inducible defensive structures when predators are present (Miyakawa et al. 2015). At the molecular level, plasticity is achieved through mechanisms such as alternative splicing, which enables the production of multiple protein isoforms from a single gene and allows cells to adapt to changing conditions (Biamonti et al. 2019). A compelling example of environmentally triggered molecular plasticity is the pea aphid, which produces winged or wingless offspring depending on population density and food availability, a switch governed by ecdysone signaling (Vellichirammal et al. 2017). Together, these examples illustrate a broad spectrum of plasticity that becomes increasingly complex across the animal kingdom, spanning temporal scales from rapid, reversible molecular responses to organismal remodeling over the course of development.

Animal pigmentation represents a particularly instructive model of phenotypic plasticity, as color change spans a continuum from rapid physiological responses to slow morphological changes, and serves diverse adaptive functions, including camouflage, predator avoidance, and intraspecific communication (Stuart-Fox and Moussalli 2009; Stevens 2016). These mechanisms can be broadly categorized into three types. Neural color change is the fastest, occurring within milliseconds to seconds, and is mediated by neuroendocrine or neuromuscular control; in cephalopods, for instance, radial muscle activation rapidly adjusts chromatophore size (Laan 2014), while octopamine and serotonin modulate both the speed and the patterning of color change (Guan et al. 2017). Physiological color change operates over minutes to hours via receptor-mediated hormonal signals. In harlequin poison frogs, coordinated action of α-MSH and MCH controls coloration (Posso-Terranova and Andrés 2017), and in barn owls, glucocorticoid receptor expression covaries with stress-related pigmentation signals (Béziers et al. 2019). While neural and physiological mechanisms enable rapid, reversible adjustments, they operate within a pre-existing cellular landscape. Morphological color change, by altering the number, type, and distribution of pigment cells, may set the functional boundaries (or dynamic range) over which fast color change can occur. As a result, morphological color changes unfold over days to weeks through the synthesis, redistribution, or migration of pigment cells — as seen in the ontogenetic skin color changes of American alligators, which involve pigment cell migration rather than changes in pigment concentration within existing cells (Stevens 2016; Francis et al. 2023). Despite this diversity, research on animal coloration (particularly in cephalopods) has disproportionately emphasized rapid neural mechanisms, often treating them as the primary drivers of color change while overlooking slower, substrate-level processes that may fundamentally shape these responses.

Cephalopods have become a compelling system for studying dynamic color change owing to their elaborate chromatophore arrays and remarkable camouflage capabilities (Sutherland et al. 2008). However, research on this system has been overwhelmingly biased toward neural control of chromatophore expansion, emphasizing millisecond-scale responses while largely overlooking the slower processes that establish the pigmentary substrate. As a result, it remains unclear whether cephalopods exhibit meaningful morphological color change through alterations in pigment ell density, composition, or distribution, or how such changes may constrain or extend the range of neural responses. Notably, most squid species are pelagic, limiting opportunities for direct observation and controlled study (Nakajima et al. 2022). *Sepioteuthis lessoniana*, the bigfin reef squid, is an exception. Distributed across the Indian Ocean and western-central Pacific near shores and coral reefs (Izuka et al. 1994; Satjarak et al. 2021), it is semi-pelagic across its life cycle. It is typically found in midwater but also displays bottom-associated and benthic behavior throughout its lifespan (Nakajima et al. 2022). Juveniles are primarily in the water column, while adults occupy a broader range, shifting to a semi-benthic lifestyle and, accordingly, to a different coloration. This ontogenetic and ecological transition provides a natural framework to test whether shifts in environmental context are accompanied not only by changes in neural patterning but also by remodeling of the pigmentary substrate underlying these responses.

Here, we ask whether morphological color change contributes to the phenotypic plasticity in a cephalopod system traditionally viewed as dominated by neural control, and whether such changes alter the baseline upon which rapid color responses operate. This study investigates the cellular mechanisms underlying morphological color change in *S. lessoniana* by examining dermal chromatophore units across developmental stages and rearing conditions. By rearing hatchlings in black or white environments for 14 weeks, we systematically assessed how age, environment, and body region (dorsal vs. ventral) interacted to influence the density and expansion of black, yellow, and red chromatophores. We utilized high-resolution brightfield microscopy and automated image analysis to track these shifts, allowing us to identify physiological trade-offs between pigment cell quantity and size. To our knowledge, this is the first study to assess basal differences in cephalopod color change at the cellular level across development, providing critical insights into the timing and limits of phenotypic plasticity in a highly adaptive marine invertebrate. By linking environmental conditions to pigment cell architecture across development, this study tests whether slower structural forms of plasticity recalibrate the dynamic range of neural color change, providing a more integrated view of how phenotypic plasticity operates across timescales.

## METHODS

### Animal husbandry

Bigfin reef squid, *Sepioteuthis lessoniana* (Shiro-ika), were hatched at the Okinawa Marine Science Station (OMSS) at Okinawa Institute of Science and Technology (OIST) Graduate University. Egg strings containing 9 eggs or more (attached to seagrass beds along the west coast of Okinawa, Tancha Village, 26.5058,127.87) were collected in March 2023 (Segawa et al. 1993; Izuka et al. 1994; Izuka et al. 1996). Tanks were illuminated by fluorescent lighting on a 12-hour day/night cycle and supplied with ocean water via an open-flow-through system. In the first month, animals were fed 3-4 times daily with a combination of live mysid shrimps and dead fish larvae and shrimp. After this, live food was eliminated from the diet (Satjarak et al. 2021). Dead squid, waste, and any remaining food were removed from the tank after feeding sessions or upon discovery (Nakajima et al. 2022).

### Animal rearing, tissue collection, and microscopy

Hatchlings were collected from the incubation tank using a clear plastic container and distributed into six glass rearing tanks. Four tanks (50 × 40 × 40 cm, ~60 L) were stocked with 40 hatchlings each, while two larger tanks (50 × 60 × 40 cm, ~90 L) were stocked with 60 hatchlings each. Tanks were covered on all sides and bottom with black or white panels to create the dark and light environments. Tanks were enclosed on 5 sides, with the top open. *S. lessoniana* were anesthetized in 4% ethanol solution and sacrificed via pithing at 4 (n= 60), 9 (n=27), and 14 weeks (n=14). Animals were collected at random and not sexed; we assume an equal distribution of males and females. Whole mantles were dissected and fixed in 4% paraformaldehyde. 4 mm diameter hole punches of the dorsal and ventral sides of the mantle were taken, then mounted on 75mm × 25mm microscopy slides for brightfield microscopy. Slides were imaged using a Keyence BZ-X800. The imaging depth varied with section thickness and was taken when chromatophores were in focus. Images were captured in sections for maximum clarity and stitched before image analysis. Each punch was imaged in 4 sections, then stitched.

### Image analysis

Images were calibrated (4×4 mm, 3000×3000 pixels) and analyzed using ImageJ (Schindelin et al. 2012). To standardize our work, we developed a macro script to classify three types of chromatophores (dark, red, and yellow chromatophores) and to quantify the total number and area of all punches collected. Chromatophores were classified by size, color, and texture (Williams et al. 2019; Bower et al. 2024) (Figure 1). At the 4-week time point, chromatophores that were not dark pigment-containing chromatophores were collectively classified as yellow pigment chromatophores because the distinction between red- and yellow-pigment chromatophores is not clearly discernible. Images were excluded if they showed no chromatophores due to bleaching or mishandling during preparation (n= 18).

**Figure 1.**
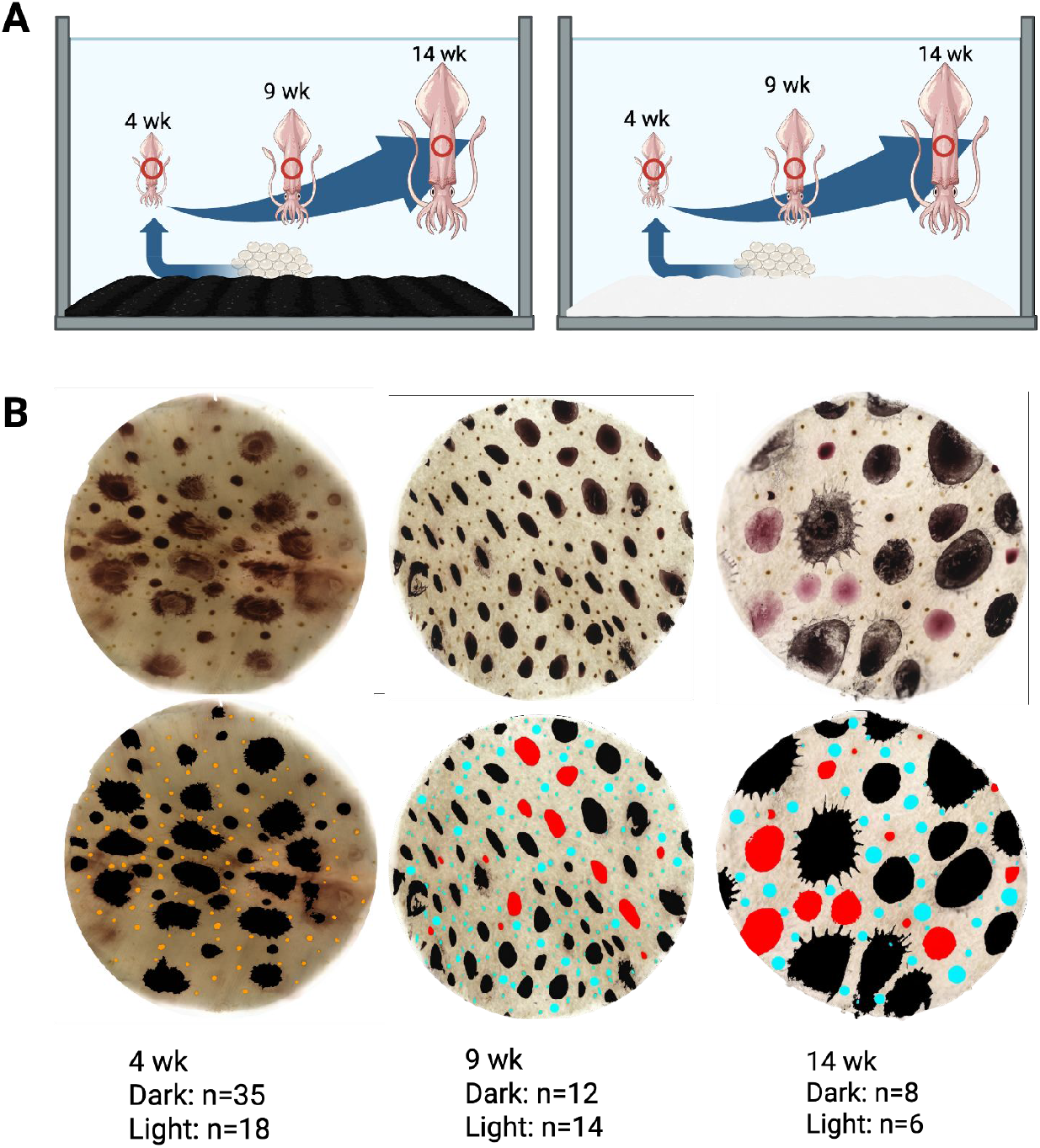
Experimental design for behavior analysis. (A) Tank setup. Red circles indicate region where punches were taken from on the dorsal and ventral mantle. (B) Example tissue punches from squid at each timepoint post-hatching before (top) and after (bottom) quantification and false-color coding of black, red, blue for yellow-pigment cells and combined red/yellow (orange).

### Statistical Analyses

Levene’s test was used to assess homogeneity of variance across the datasets; the test was significant for all datasets, indicating high variance among the observations. To account for nonindependence of observations, the data were analyzed using a weighted Linear Mixed Model (LME), treating individual subject IDs as a random effect. To maintain the integrity of the error estimates, the model was adjusted with variance structures designed to mitigate heteroscedasticity associated with age and background variables. Next, a 2×3×2 mixed-effects analysis of variance (ANOVA) was employed to assess the impact of Environment (white/black), Age (four-, nine-, and fourteen-week-old), and Body Region (dorsal/ventral) on chromatophore density and distribution. And finally, post hoc pairwise comparisons were performed using Tukey’s Honestly Significant Difference (HSD) test to identify specific group differences. All analyses were conducted in R (Team 2025) (version 4.3.1) with a statistical significance set at α = 0.05. Area values were normalized by dividing the total area by the chromatophore number per individual

## RESULTS

### Morphological Dynamics of Black Pigment Dispersal: Developmental Shifts in Cell Size and Area

A weighted mixed-effects ANOVA showed that Environment and the Age × Body Region interaction significantly affect black pigment cell count. This analysis showed a significant Age x Body Region interaction (F_2, 92_ = 14.67, p <0.0001) on count. Post-hoc analysis showed that at every age, the dorsal side (Δ = 11.3, SE = 1.58, p < 0.0001) had significantly more black-pigment cells (Figure 2A). As the animal grows, the difference between the dorsal and ventral side increases, with the largest difference occurring at 14 weeks (Δ = 18.77, SE = 3.39, p <0.0001). This is mainly due to a sharp decline in the number of black-pigment cells on the ventral side with age. There was also a significant main effect of the rearing environment (F = 40.01, p < .0001). Across all groups, black-reared individuals had a higher number of black-pigment cells than white-reared individuals (Δ = 0.37, SE = 1.73, p < 0.0001; Figure 2B).

**Figure 2.**
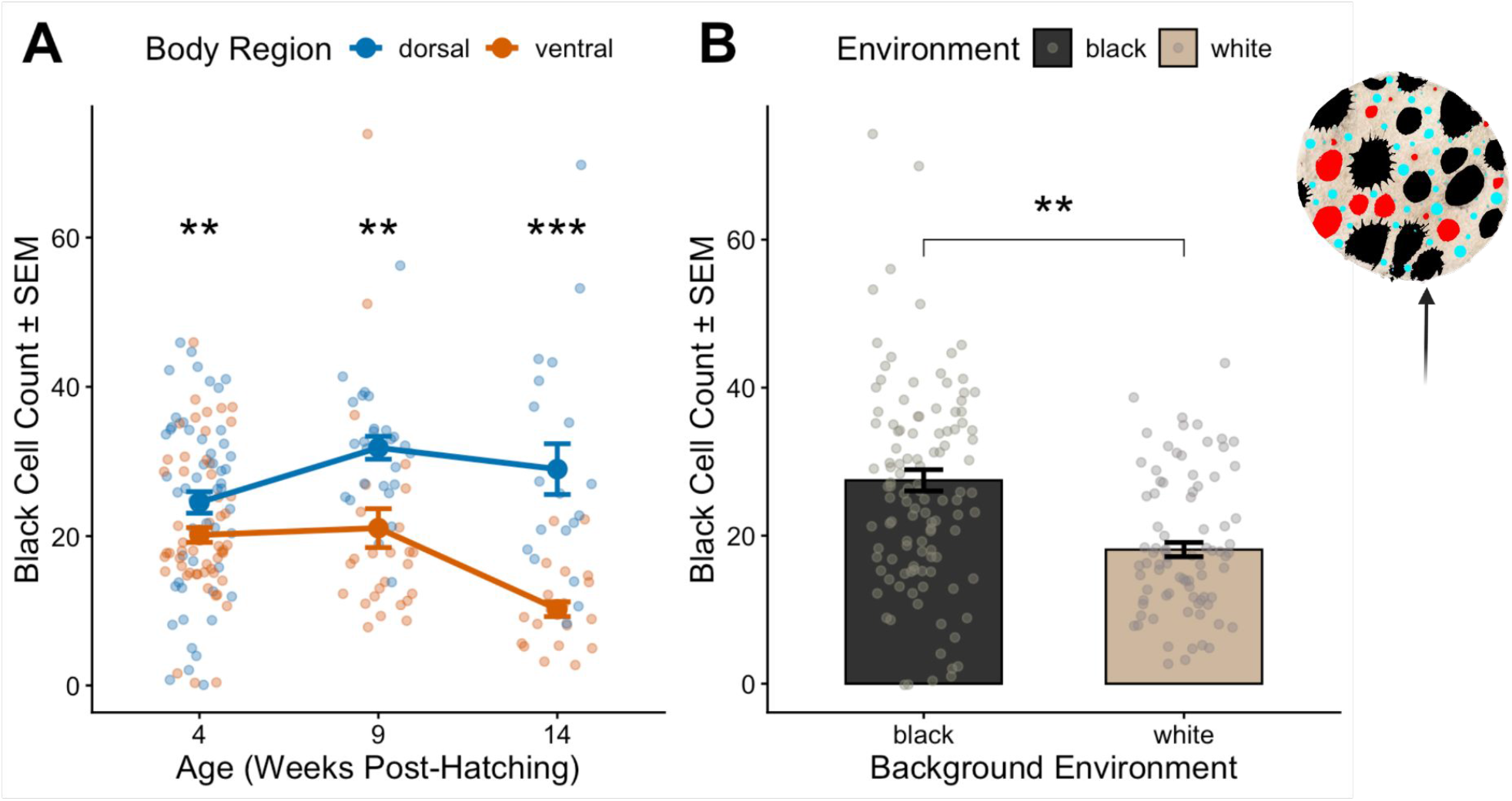
Developmental and environmental change of black-pigment cell quantity. (A) Mean black-pigment cell counts (±SEM) across developmental stages (4, 9, and 14 weeks post-hatching), separated by Body Region: Dorsal (blue) and Ventral (orange), n=196 (98 dorsal, 98 ventral). (B) Main effect of background environment. Mean black-pigment cell counts (averaged across Age and Body Region) for black-reared (black) and white-reared (white) individuals. Analysis was conducted using a weighted Linear Mixed Model (LME) with individual ID as a random effect and variance structures to account for heteroscedasticity across age and background groups. n=196 (82 white, 114 black). *p < 0.05, **p < 0.01, ***p < 0.001

When examining the dispersal of black-pigment cells, we found a significant Age × Body Region interaction (F(2, 134) = 8.39, p < 0.001). Early in development (Weeks 4 and 9), the dorsal pigment cells are significantly larger than the ventral (Δ = 0.03, SE = 0.004, p <0.0001), however, by the last timepoint, this pattern switches, and the ventral pigment cells are significantly larger (Δ = 0.069, SE = 0.045, p <0.05; Figure 3A). The few black-pigment cells that remain on the ventral side show increased expansion, and therefore, each ventral cell contributes more toward coloration than those on the dorsal side. We also observe a significant Environment × Age interaction (F(2, 134) = 12.47, p < 0.0001). At 4 weeks, the black-reared animals have larger black-pigment cells (Δ = 0.012, SE = 0.0051, p < 0.05), but this quickly reverses at 9 weeks, at which point the white-reared animals have significantly larger cells (Δ = 0.047, SE = 0.013, p < 0.001). Although not significant, this trend continues to 14 weeks. The white-reared squid have much wider variation in cell size than black-reared animals at this time point (Figure 3B). There appears to be a trade-off regarding the quantity of black-pigment cells and size.

**Figure 3.**
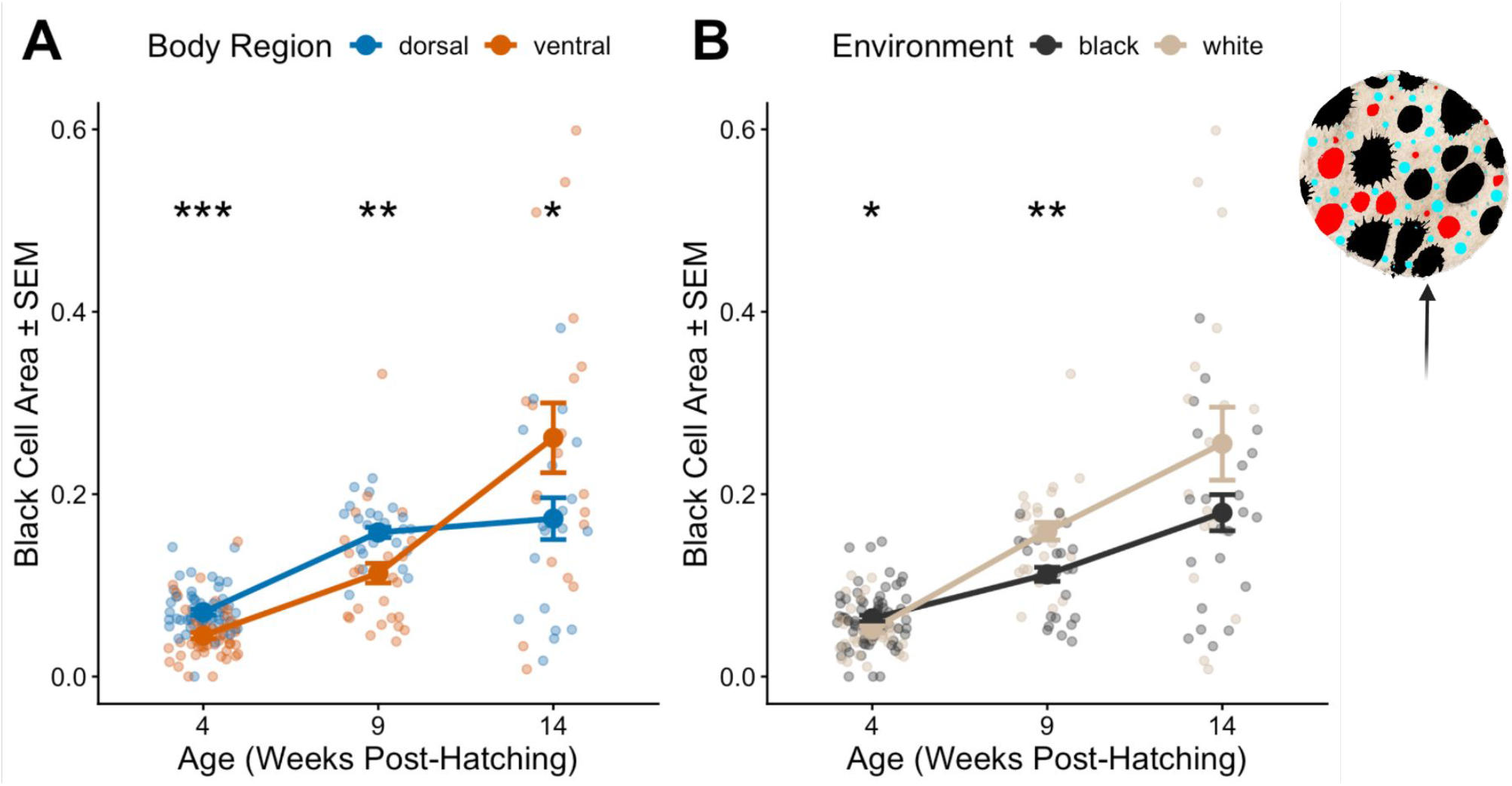
Developmental and environmental change of black-pigment cell area. (A) Mean black-pigment cell area (±SEM) across developmental stages (4, 9, and 14 weeks post-hatching), separated by body region: dorsal (blue) and ventral (orange). n= 198 (99 dorsal, 99 ventral) (B) Main effect of background environment. Mean black-pigment cell area (±SEM) across developmental stages (4, 9, and 14 weeks post-hatching), separated by environment: black-reared (black) and white-reared (white) individuals n= 198 (82 white, 116 black). Analysis was conducted using a weighted Linear Mixed Model (LME) with individual ID as a random effect and variance structures to account for heteroscedasticity across age and background groups (*p < 0.05, **p < 0.01, ***p < 0.001)

### Yellow Chromatophore Proliferation: Black-Reared Squid Exhibit a Mid-Developmental Surge in Cell Count

We observe significant effects on the number of yellow-pigment cells for the Age x Environment interaction (F(2,92) = 6.24, p < 0.05) and the Age x Body Region interaction (F_2,92_ = 14.33, p < 0.001). Overall, the black-reared squid have a higher number of yellow-pigment cells than white-reared squid at 9 (Δ = 51.08, SE = 11.8, p <0.0001) and 14 weeks (Δ = 37.56, SE = 12.0, p<0.05). The black-reared squid had a massive increase in cell number at 9 weeks, whereas cell quantity in the white-reared group remained stable throughout development (Figure 4A). Comparisons between Body Regions at each developmental stage remain similar until later in development. Body regions showed no significant differences in cell number until the 14-week mark, at which point ventral pigment cells significantly decreased (Δ=34.96, SE = 8.37, p<0.0001; Figure 4B).

**Figure 4.**
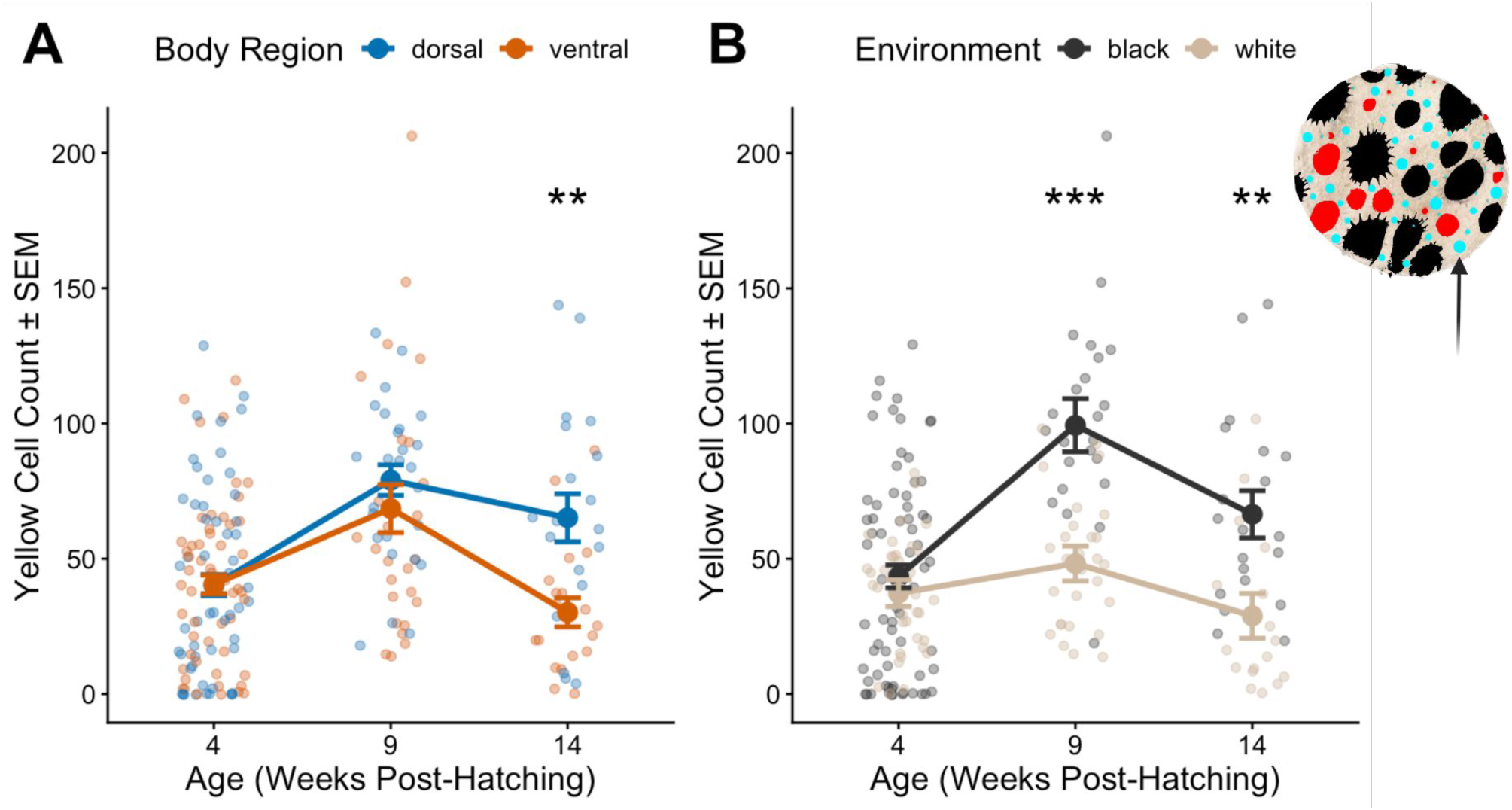
Developmental and environmental change of yellow-pigment cell quantity.(A) Mean yellow-pigment cell count (±SEM) across developmental stages (4, 9, and 14 weeks post-hatching), separated by environment: black-reared (black) and white-reared (white) individuals. **n= 196 (82 white, 114 black).** B) Mean yellow-pigment cell count (±SEM) across developmental stages (4, 9, and 14 weeks post-hatching), separated by body region: dorsal (blue) and ventral (orange). **n= 196(98 dorsal, 98 ventral)** Analysis was conducted using a weighted Linear Mixed Model (LME) with individual ID as a random effect and variance structures to account for heteroscedasticity across age and background groups (*p < 0.05, **p < 0.01, ***p < 0.001)

There is a significant three-way interaction among Age, Environment, and Side (F (2, 134) = 4.82, p < 0.01) for yellow-pigment cell area. On the ventral side, the pigment area is highly sensitive to age and rearing environment. Black-reared squid have significantly larger cells at 4 weeks (Δ = 2.06×10^−3^, SE = 0.0006, p<0.001), but they decrease in size with age. In contrast, the pigment cell area increases with age in the white-reared squid and, by the last timepoint, they have larger cells than the black-reared group, although this remains extremely variable (Figure 5). On the dorsal side, the cell area is much more stable across both rearing environments. There is a significant difference in early development (Δ = 8.58×10^−4^, SE = 0.0003, p<0.01), but this trend does not continue with age (Figure 5).

**Figure 5.**
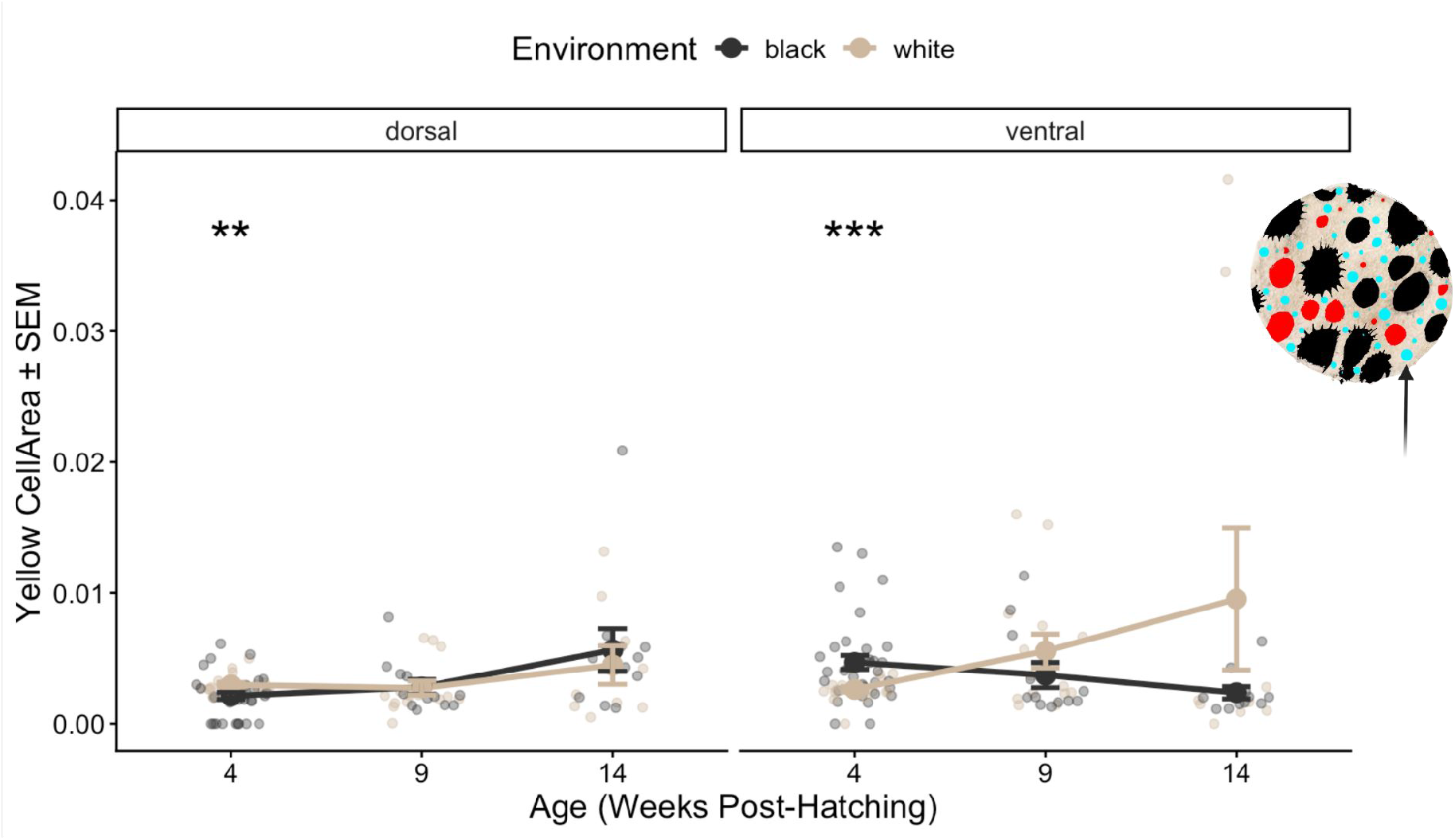
Yellow-pigment cells show plasticity specifically from white-reared on the ventral side. Mean yellow-pigment cell area (±SEM) across developmental stages (4, 9, and 14 weeks post-hatching), separated by environment: black-reared (black) and white-reared (white) individuals and region. **n= 198 (82 white, 116 black).** Analysis was conducted using a weighted Linear Mixed Model (LME) with individual ID as a random effect and variance structures to account for heteroscedasticity across age and background groups (*p < 0.05, **p < 0.01, ***p < 0.00).

### Interactive Effects of Age and Environment on Red-Pigment Cell Area

There is a significant effect of age on red-pigment cell number (F_1,41_ = 12.29, p < 0.001), with significantly more pigment cells at 14 weeks compared to 9 weeks (Δ = 15.4, SE = 4.35, p<0.01; Figure 6). Unlike other pigment cell types, red cell number is not affected by region or rearing conditions. However, red-pigment cell area shows significant differences for Age x Environment interaction (F_1,49_ = 13.50, p < 0.01) and Body Region (F_1,49_ = 92.05, p < 0.0001).

**Figure 6.**
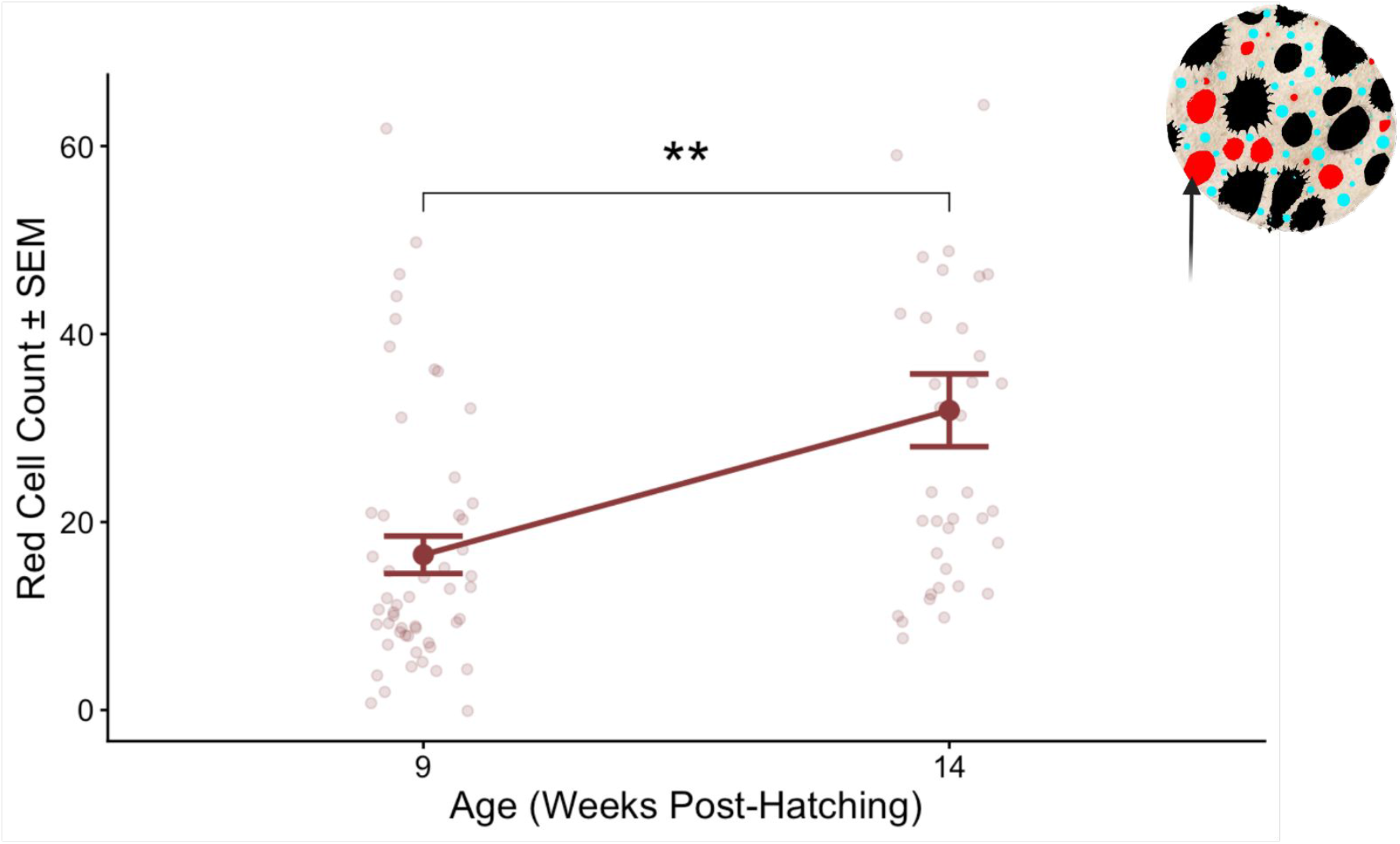
Red-pigment cell numbers increase with age (A) Mean red-pigment cell count (±SEM) across developmental stage averaged across Environment and Body Region n= 90 (52: 9 weeks, 38:14 weeks). Statistical analysis was conducted using a weighted Linear Mixed Model (LME) with individual ID as a random effect and variance structures to account for heteroscedasticity across age and background groups (*p < 0.05, **p < 0.01, ***p < 0.00)

The black-reared individuals do not change in size with age. In contrast, the white-reared individuals show an increase in cell size between 9 and 14 weeks. Specifically, at 9 weeks, the white-reared individuals have significantly smaller cells than the black-reared individuals (Δ = 0.011, SE = 0.0053, p<0.05), but by 14 weeks, they have larger red-pigment cells (Δ = 0.024, SE = 0.013, p=0.063; Figure 7). Similar to the other pigment cell types, the red-pigment cells in the ventral region are significantly larger than in the dorsal region (Δ = 0.033, SE = 0.0067, p<0.001; Figure 7B). At the 4-week timepoint, the combination of small cells (yellow and red pigment cells) is significantly different between rearing conditions on the dorsal (Δ = 0.00086, SE = 0.00032, p<0.05) and ventral (Δ = 0.002061, SE = 0.000581, p<0.001; Figure 8) sides.

**Figure 7.**
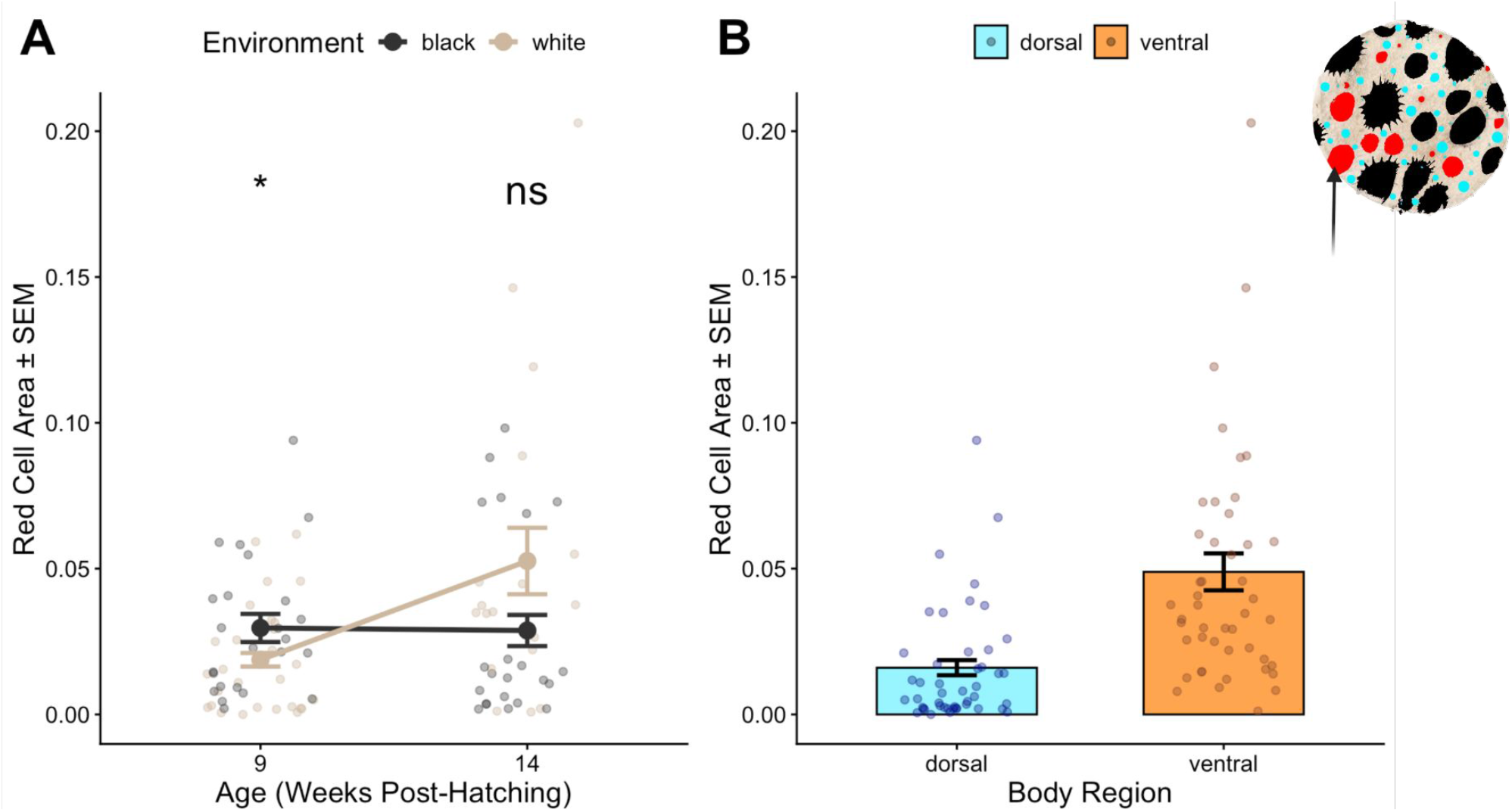
Red-pigment cells are larger on the ventral side for white-reared individuals (A) Main effect of background environment. Mean red-pigment cell area (±SEM) across developmental stages (4, 9, and 14 weeks post-hatching), separated by environment: black-reared (black) and white-reared (white) individuals **n= 92 (46 white, 46 black).** (B) Mean red-pigment cell area (averaged across Age and Environment) separated by body region:dorsal (blue) and ventral (orange) **n= 92 (46 dorsal, 46 ventral)**.Statistical analysis was conducted using a weighted Linear Mixed Model (LME) with individual ID as a random effect and variance structures to account for heteroscedasticity across age and background groups (*p < 0.05, **p < 0.01, ***p < 0.00).

**Figure 8.**
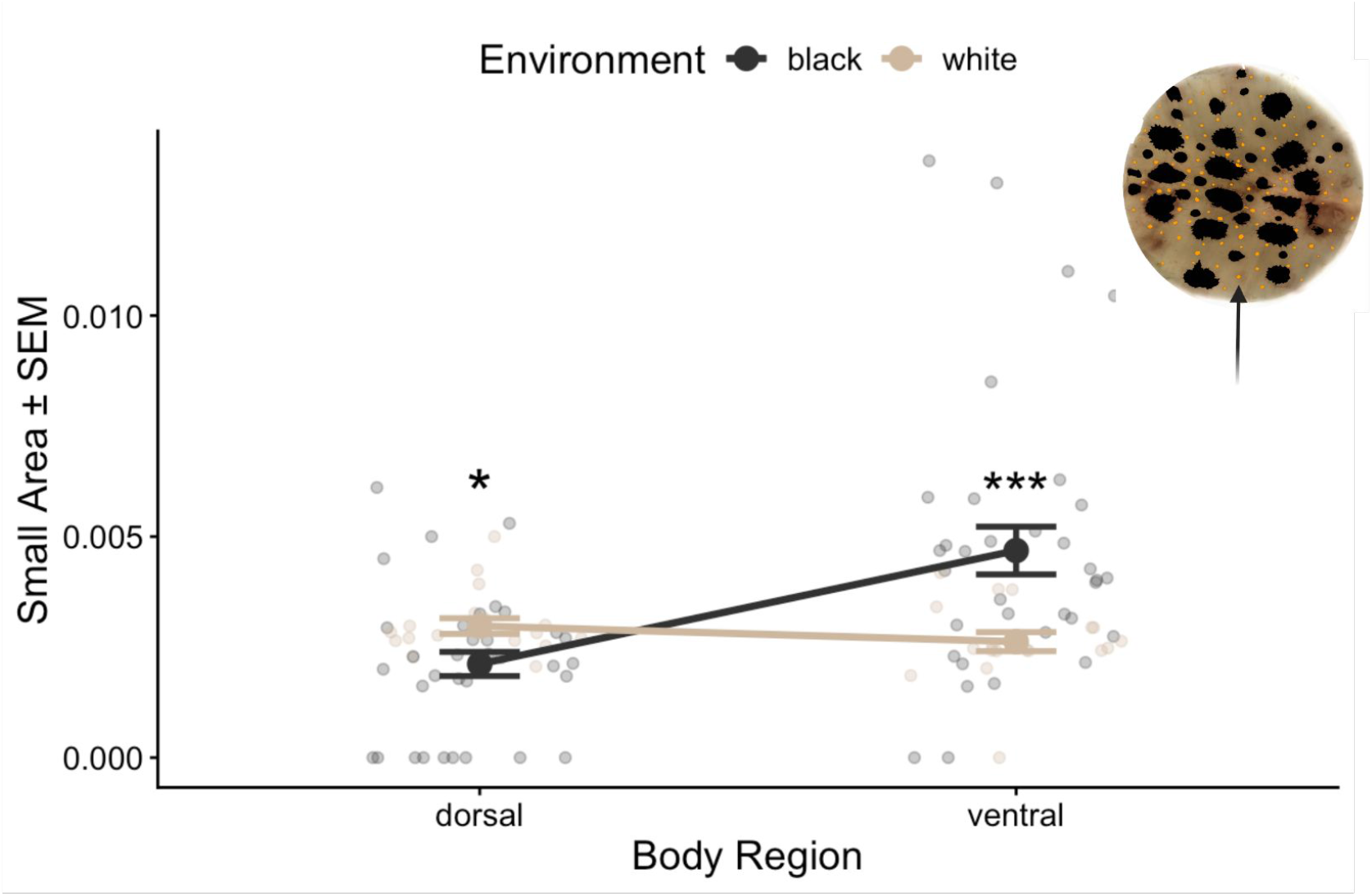
Red and yellow pigment cells are larger on the ventral side at a young age (A) Main effect of background environment. Mean small pigment cell area (±SEM) at week 4 on Body Region separated by environment: black-reared (black) and white-reared (white) individuals n= 106 (36 white, 70 black). Statistical analysis was conducted using a weighted Linear Mixed Model (LME) with individual ID as a random effect and variance structures to account for heteroscedasticity across age and background groups (*p < 0.05, **p < 0.01, ***p < 0.001)

## DISCUSSION

The results of this study demonstrate that *Sepioteuthis lessoniana* employs an age-dependent, environmentally responsive strategy for morphological color change. Dermal pigmentation is not fixed, but shifts dynamically across development in response to background reflectance. We identify a clear trade-off in chromatophore organization: individuals either maintain a high density of small chromatophores or a lower density of larger, more expansive ones. A primary finding was the significant interaction between environment and age regarding black-pigment chromatophore cells. In black-reared environments, squid maintained a higher total dark pigment chromatophore count. Conversely, white-reared squid exhibited a distinct strategy: while they had fewer black cells, those cells were significantly larger at the 9- and 14-week marks. This suggests that when the energetic cost of maintaining a high cell density is not supported by the environment, the animal may maximize the efficiency of existing cells through increased dispersal or muscular expansion.

The sharp decline of dark chromatophores on the ventral side with age, coupled with the increasing difference between dorsal and ventral counts, aligns with the development of organismal countershading. As *S. lessoniana* matures and moves into more open-water environments, the evolutionary pressure to remain dark on top (to blend with the depths) and light on the bottom (to blend with the surface light) becomes more pronounced. Interestingly, the few dark pigment chromatophores that remain on the ventral side at 14 weeks are significantly larger than those on the dorsal side. This developmental trajectory provides evidence that phenotypic plasticity in color change is not a static capacity but is calibrated over time through experience and maturation, situating *S. lessoniana* at the intersection of behavioral, physiological, and morphological plasticity in ways that few other organisms permit. This pattern suggests that countershading is achieved not only through differential chromatophore distribution but also through differential expansion capacity, with ventral regions relying on fewer but more expansive chromatophores to maintain effective light matching.

The ontogenetic shift from a predominantly pelagic juvenile stage to a more structurally complex, semi-benthic adult niche provides a compelling ecological context for these changes(Vidal and Shea 2023). Early-life environments in the water column likely favor uniform, low-contrast body patterns, whereas later transitions to reef-associated or benthic habitats require increased spatial and chromatic complexity(Chung and Marshall 2016). Our data suggest that chromatophore architecture is developmentally tuned to these shifting ecological demands, effectively preconfiguring the substrate upon which well-known rapid neural patterning operates. In this sense, morphological color change may function as a form of developmental “priors,” constraining and enabling the range of neural outputs available to the animal as it transitions across ecological niches. Changes in chromatophore density and size likely alter the effective “color space” available to the organism(How et al. 2017). High-density configurations may permit fine-grained patterning but limit maximal contrast, whereas low-density, large-cell configurations may enable greater contrast at the expense of spatial resolution. The observed density–size trade-off suggests that squid dynamically tune the resolution and amplitude of their color output in response to environmental context.

Cephalopod color change has historically been conceptualized as a primarily neural phenomenon, with chromatophore expansion treated as the dominant axis of variation(Mattiello et al. 2010; Hanlon and Messenger 2018). However, our findings challenge this view by demonstrating that the cellular substrate itself is dynamically remodeled over developmental time. This suggests that neural control operates on a shifting morphological landscape rather than a fixed one, and that the limits of pattern generation may be fundamentally constrained by chromatophore density, size, and spatial distribution (Alvarado 2020). Importantly, these findings reinforce the idea that phenotypic plasticity in cephalopods operates across multiple temporal scales. While neural mechanisms enable rapid, reversible changes on the order of milliseconds, our results demonstrate that the underlying pigmentary substrate is itself plastic over weeks, effectively recalibrating the baseline from which rapid responses are deployed.

Together, these findings support a hierarchical model of color change in cephalopods, in which slow morphological plasticity defines the bounds of a rapidly deployable neural system. To our knowledge, we provide the first cellular-level evidence of age-restricted phenotypic plasticity in *S. lessoniana*. The discovery of a density-size trade-off and the characterization of the 9-week developmental surge offer new insights into how cephalopods optimize pigment expression to survive in fluctuating environments. Future research should investigate the metabolic costs of maintaining high chromatophore counts relative to the muscular energy required for sustained cell expansion. These alternative strategies likely reflect distinct energetic regimes. Maintaining a high density of chromatophores may entail developmental and metabolic costs associated with pigment synthesis and cellular maintenance, whereas relying on fewer, larger chromatophores may shift the burden toward the neuromuscular control required for sustained expansion. This suggests that chromatophore organization may represent an optimization problem that balances developmental investment with real-time performance costs. Together, these findings suggest that cephalopod color change is not solely a product of rapid neural control, but rather emerges from the interaction between fast physiological processes and slower, developmentally tuned morphological substrates that define the boundaries of adaptive visual signaling.

